# Phylogenies and diversification rates: variance cannot be ignored

**DOI:** 10.1101/406504

**Authors:** Daniel L. Rabosky

## Abstract

The concept of variance is the foundation of modern statistics; it reflects our awareness that independent samples from a single population or stochastic process can produce a range of outcomes. A recent pair of articles in the journal *Evolution* abandons the notion of the sample variance and advocates for uncorrected comparisons of numerical point estimates between groups. The articles in question (Meyer and Wiens 2017; Meyer *et al*., 2018) criticize BAMM, a scientific software program that uses a Bayesian mixture model to estimate rates of evolution from phylogenetic trees. The authors use BAMM to estimate rates from large phylogenies (n *>* 60 tips) and they apply the method separately to subclades within those phylogenies (median size: n = 3 tips); they find that point estimates of rates differ between these levels and conclude that the method is flawed, but they do not test whether the observed differences are statistically meaningful. There is no consideration of sampling variation and its impact at any level of their analysis. Here, I show that numerical differences across groups that they report are fully explained by high variance in their subclade estimates, which is approximately 55 times greater than the corresponding variance for estimates from large phylogenies. Variance in evolutionary rate estimates – from BAMM and all other methods – is an inverse function of clade size; this variance is extreme for clades with 5 or fewer tips (e.g., 70% of clades in the focal study). The articles in question rely on negative results that are easily explained by low statistical power to reject their preferred null hypothesis, and this low power is a trivial consequence of high variance in their point estimates. By ignoring variance, the testing approach outlined in these articles can be misused to demonstrate that all statistical estimators, including the arithmetic mean, are “flawed”. I describe additional mathematical and statistical mistakes that render the proposed testing framework invalid on first principles. Evolutionary rates are no different than any other population parameters we might wish to estimate, and biologists should use the training and tools already at their disposal to avoid erroneous results that follow from the neglect of variance.

## Introduction

The concept of sampling variation is at the heart of modern statistical practice. Fundamentally, terms that describe variation—especially the notions of standard deviation and variance—encapsulate the idea that independent samples from a single population are unlikely to be identical (Pearson 1896, Galton 1907, Fisher 1918, Fisher 1925). It is the variance in the sampling distribution that characterizes how different these samples might be. Suppose I wish to determine whether two populations share a common mean, given that I have obtained samples from each population with respective sample means of 5 and 50. It is trivially true that 5 is less than 50, and yet we can draw no conclusions whatsoever about the meaning of this difference without an estimate of the sample variance. If the variance is high, repeated sampling of the first population may be just as likely to produce a little number (“5”) as a large number (“50”). In similar fashion, independent samples (observations) from a single stochastic process can generate a range of outcomes, even if the underlying process is identical. For example, we generally believe that the process of nucleotide substitution is effectively stochastic when considered over phylogenetic timescales. Hence, we find it unsurprising that some lineages will, just by chance, accumulate more (or fewer) DNA sequence substitutions relative to other lineages, over a given amount of time. There are very few measurable quantities in science – perhaps none – that are truly independent of sampling variation. For more than two centuries, scientists have recognized a need to account for sampling variation when comparing numerical point estimates (e.g., means) between populations: the use of probability to characterize the variation in sample outcomes can be traced back to Laplace (late 1700s) or perhaps even earlier (Stigler 1973, Stigler 1986).

Variation in independent estimates of a population parameter can result from at least three general causes. First, variation might reflect the stochastic outcomes of a sampling process that estimates a parameter of interest from a finite number of observations (e.g., how many heads in a sample of 5 coin tosses?). Second, variation can reflect a complex intersection of multiple and potentially unmeasurable deterministic events; we may find it useful or even necessary to conceptualize the net effect of these events as a stochastic process. For example, even if DNA sequence evolution is fully deterministic at the lowest level of biological organization, it is nonetheless useful to approximate the outcome of these deterministic processes using a stochastic Markov chain model. Indeed, this approximation is the foundation of the entire field of molecular evolution and is essential for modern phylogenetic tree inference. Finally, variation might reflect measurement error from the instruments we use to gather information about the physical and biological properties of the world in which we live. Throughout the remainder of this article, I use the catchphrase “sampling variation” to encapsulate variation in a sample that might be attributable to any of these three factors.

We live in a world in which virtually all published research entails attention to sampling variation on some level. Against this backdrop of general statistical rigor, two recent articles published in Evolution have used numerical point estimates, without characterizing sampling variation, to produce sweeping generalizations about model adequacy. These articles (MW2017: Meyer and Wiens 2017; MRW2018: Meyer et al. 2018; corresponding author J. J. Wiens on both articles) have openly eschewed standard practice with respect to characterizing the impact of sampling variation, in that such variation is ignored altogether. Both articles are critiques of BAMM, a software program developed by my research group to estimate evolutionary rates from phylogenetic trees. There are numerous issues to be resolved with BAMM and other approaches for modeling evolutionary rates on phylogenies, including the adequacy of the assumed likelihood function and prior model (Moore et al 2016; Rabosky et al. 2017), and I welcome research attention to identify solutions to these and other issues. However, the general statistical errors in MW2017 and MRW2018 have little to do with BAMM. As I demonstrate below, their results are the predictable outcome of ignoring sampling variation in favor of raw numerical point estimates that are highly influenced by such variation. Most worryingly, the authors exhort other researchers to perform similar analyses on their data and to use these tests as a basis for rejecting the use of hierarchical models more generally. By ignoring sampling variation and its consequences, the analyses espoused by MW2017 and MRW2018 are deeply flawed.

There are numerous errors of statistics, mathematics, and logic in MW2017 and MRW2018, and I address the most significant of these in considerable detail. First, I describe the precise comparison advocated by MW2017 and MRW2018 and explain why their approach represents both a statistical and mathematical error. I then illustrate that all statistical estimators, including the arithmetic mean, can be shown to fail when sampling variation is ignored. I then demonstrate exactly how MRW2018 obtained the results they present, and I show that their findings are fully predictable from massive and unaccommodated sampling variation that resulted from their analysis of datasets (phylogenies) that contained fewer than 5 tips. By estimating “population means” using a median sample size of merely 3 individuals (observations; data points) per population, the variance associated with their estimates is far beyond what would is typically considered acceptable in any scientific discipline. Finally, I describe several theoretical errors that render the testing procedure of MW2017 and MRW2018 invalid on first principles.

## Scope of the present article

**There are only two analyses in MRW2018 that bear on the validity of BAMM; they are described in the MRW2018 section titled *“BAMM gives misleading results in empirical studies”*.** My analysis in this article bears strictly on the testing framework described and applied in that section of their article. The simulation-based analyses from MW2017 contending that BAMM is flawed were decisively addressed by Rabosky (2018) and will not be revisited here. The reader of this thread of articles is warned that much of the text in MW2017 and MRW2018 has no bearing on BAMM but is presented as advocacy for an alternative inference framework. For the present purposes, I am not comparing BAMM to alternative inference frameworks, and I am not providing new tests with respect to whether BAMM does or does not work. The present article and Rabosky (2018) are strictly intended to describe mathematical and statistical errors in the testing procedures advocated by MW2017 and MRW2018. R code to reproduce all analyses described here is provided on Github (github.com/macroevolution/variance). The relevant scripts and data files are linked at the end of each section below.

## 1. Comparative framework in MW2017 and MRW2018 is not valid

The empirical test in MW2017 and MRW2018 involves the comparison of evolutionary rates for groups (clades) derived from two non-identical datasets. If the point estimates differ, the authors conclude that the method is flawed. The datasets used to obtain the point estimates differ in that one is a nested subset of the other. I label the two estimation strategies from MRW2018 as “use-all-data” and “discard-data” throughout, because one approach involves performing an analysis with a greatly reduced subset of the full dataset (e.g., a taxonomic genus-level versus family-level analysis). These approaches can be described as follows (Fig. 1):

**Figure 1.**
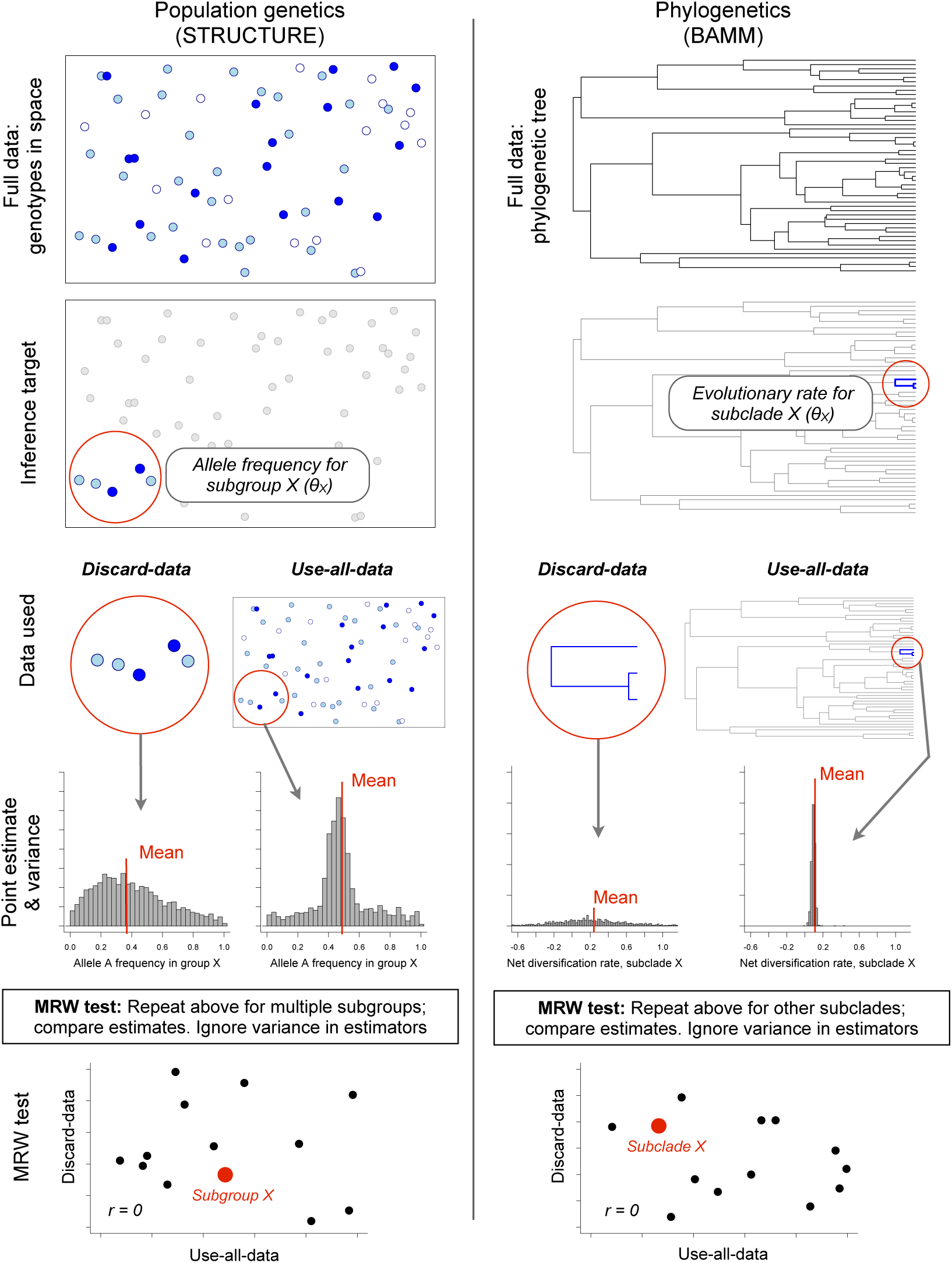
Illustration of testing procedure advocated by MW2017 and MRW2018, illustrated for Bayesian clustering methods in population genetics (program STRUCTURE) and phylogenetics (BAMM). Full data for STRUCTURE is a set of genotypes arrayed in space (1-locus example shown here: dark blue = AA; light blue = Aa; white = aa); full data for BAMM is a phylogenetic tree. Estimation of parameters is then performed for a targeted subgroup of the data. For STRUCTURE, the inference target might be allele frequencies in a local cluster of individual genotypes; for BAMM, the target might be evolutionary rates for a single genus with 3 species (blue). Estimates are then made using two strategies, referred to as “discard-data” and “use-all-data”. In the “discard-data” approach, a purely local estimate of parameters is made, ignoring all information from the broader population from which the local subgroup (or genus) was drawn. For the “use-all-data” approach, the full population data (STRUCTURE) or phylogenetic tree (BAMM) is used to estimate parameters across the complete data; the marginal distribution of allele frequencies (STRUCTURE) or evolutionary rates (BAMM) is then extracted for the subgroup of interest. MRW2018 suggest that researchers should perform this analysis across multiple subgroups and compare the resulting point estimates of parameters. A non-significant correlation between parameter estimates at the use-all-data and discard-data scales for an empirical dataset is interpreted as failure of the inference method. The test thus rejects inference methods based on a negative result and is sensitive to any factors that reduce the power of the test.

### Approach 1: *use-all-data*

Apply BAMM to a large phylogeny of organisms. Extract the marginal evolutionary rates for small subclade of interest from the overall analysis. In the same way, one might apply the program STRUCTURE (Pritchard et al. 2000) to a large population genomic dataset of many individuals, then extract the marginal allele frequencies for a particular subgroup of just three individuals (Fig. 1). Similarly, one might apply a hierarchical linear model for heart disease to a nationwide health database, and then extract the marginal incidence rate for a particular subset of the full data.

### Approach 2: *discard-data*

As an alternative, BAMM is applied directly to the small subclade of interest, discarding all data from remainder of the phylogenetic dataset. In the same way, one might estimate allele frequencies by applying STRUCTURE to a reduced dataset of three individuals (Fig. 1), or one might estimate heart disease risk using data only for a particular local community or family. This approach thus discards information that might be contained in the broader phylogeny (BAMM), the population-wide distribution of allele frequencies (STRUCTURE), or in the national health database (hierarchical linear model).

Critically, MW2017 and MRW2018 state that that use-all-data and discard-data approaches should give identical or “very similar” estimates, and they claim that methods failing to agree at this level are flawed. For example, MRW2018 write: “MW introduced an approach to evaluate the accuracy of BAMM in empirical studies. They compared diversification rates estimated for individual clades when estimated across an entire tree (with many clades) relative to rates estimated for those clades in isolation. Most importantly, if the two sets of rates differ, then BAMM must be giving inaccurate rate estimates (because two different rates for the same clade cannot both be correct).”

However, this chain of inference in MW2017 and MRW2018 involves numerous errors of mathematical, statistical, and logical reasoning. A summary of the errors associated with this procedure is provided in Table 1. The authors are interested in the marginal expectation of a parameter *θ* taken over some subset of the full data *X*, or *E*(*θ*_*X*_). They compare estimates of *E*(*θ*_*X*_) from two approaches, which differ in the underlying data used to compute the estimates.

**Table 1.** Statistical, mathematical, and logical errors in the comparative test proposed by MRW2018 to determine whether a method is flawed. *Section* is the numbered subsection within the present article that describes the error

| Description of error | Type of error | Impact on MRW2018 analysis | Section |
| --- | --- | --- | --- |
| No consideration of sampling variation (variance) of estimators | Statistical | Massive inflation of variance in estimates at one level of analysis; most parameters estimated from populations (clades) with 5 or fewer observations. Variance in point estimates is sufficient to eliminate most signal from the data | 1 – 4 |
| No consideration of null hypothesis: how would test perform if all samples drawn from same true population (e.g., all clades have same true rate)? | Statistical | Test approaches 100% Type I error rate when true parameter does not vary among groups | 5 |
| MRW2018 assume that correlation coefficient is a measure of accuracy. | Statistical | Correlation is a measure of association, not of accuracy. Lack of correlation provides no information about accuracy of BAMM rate estimates, especially when applied to empirical datasets generated from unknown true rate parameters. | 6 |
| Data violate most or all assumptions of the only statistical test used in MRW2018 (Pearson correlation) | Statistical | Strong effects of outliers, nonlinearity, and heteroscedasticity. Lack of independence reduces effective number of samples to just 2-3 data points. Measures of association and significance cannot be interpreted. | 7 |
| Non-equivalence of estimators used by MRW2018. Authors use two estimators and assume they have same expected value; this assumption is false. The estimators are conditioned on different data, and likelihood equations used for inference at use-all-data and discard-data levels are different | Mathematical | Parameter estimates at use-all-data and discard-data levels are not comparable. MRW2018 test cannot be interpreted. | 1, 8 |
| Inequality of estimates is assumed to indicate absolute failure of inference method | Logical | Authors fail to consider any candidates for low correlation other than their preferred hypothesis: that the inference method is flawed. Specifically, sampling variation and all other potential causes of low statistical power are ignored. | 1 |

The first estimate of *θ*_*X*_ is conditioned only on data *X* itself; the second is conditioned on data *Y*, but where *X* is a subset of *Y* (e.g., *X∈ Y*). The use-all-data approach attempts to estimate *E*(*θ*_*X*_*| X,Y* : *X∈ Y*); the discard-data approach attempts to estimate *E*(*θ*_*X*_*| X*). Formally, MW2017 and MRW2018 make the following invalid assumption:

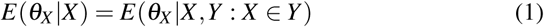

Because the two estimates are mathematically non-equivalent, the test proposed by W2017 and MRW2018 cannot be interpreted. The difference between the two estimates may or may not be of biological relevance, but the test construction itself tells us nothing without knowing the absolute accuracy of the expectations and the corresponding sampling variation. The logic error involves the subsequent chain of interpretation: the authors conclude that, if the estimates differ, the estimation method itself is flawed.

It is certainly useful to understand the conditions under which BAMM and other methods yield different results when applied to different levels of hierarchically-structured data.

But surely, if we observe that two such inference procedures appear to give different results, the most sensible and scientifically justifiable course action is to then ask: (1) How different are the estimates, in light of the observed sampling variation? (2) Is one or both set of estimates accurate in an absolute sense? (3) How would this inference procedure work if applied to another type of method, and (4) how would the comparative procedure fare if there is no variation in true population parameters (e.g., diversification rates) across the groups under consideration? None of these questions are raised or addressed by MW2017 and MRW2018. The remainder of my article dissects the consequences of neglecting these questions.

MRW2018 apply their testing procedure (Fig. 1) to two empirical datasets (tortoises and tanagers). I generally restrict my examples and reanalysis here only to the first dataset presented by MRW2018 (tortoises), because the catalog of errors I describe (Table 1) applies equally to both datasets. The authors first apply BAMM to the complete phylogeny of tortoises (63 species) and then summarize the marginal evolutionary rates for each of 13 genera from that phylogeny (“use-all-data”). They compare these point estimates to a second “discard-data” analysis where BAMM is executed separately on each of the 13 genera (median size: 3 taxa).

The authors perform no statistical assessment of whether the rates estimated using the use-all-data approach are different from those obtained using the discard-data method. The variances of the estimators are not presented, even though most of the subclades contain fewer than 5 species. The rates are not compared to true or known values, so no assessment of absolute error is possible: the authors perform this comparison only for empirical datasets where the true rates are unknown. The authors discuss the numerical differences between the sets of rates, and they perform a correlation test between the point estimates (r = 0.49, p = 0.09; n = 13 paired comparisons), which they interpret as no correlation. The point estimates are computed for empirical datasets, not for datasets simulated under known parameters; the conclusion that BAMM is flawed is derived solely from the observation that the rates are interpreted as different.

However, the use-all-data and discard-data estimators are not expected to yield the same value: they are conditioned on different data, and the simple observation that they are different tells us nothing, without knowing the variance of the estimators. The fundamental question should have involved testing a null hypothesis: is the magnitude of difference in the two sets of rates consistent with sampling variation alone? As we shall see below, sampling variation fully explains the results they obtain. Put simply, the sampling error (confidence intervals; credible intervals; “noise”) associated with each clade from the discard-data analysis is approximately 55 times greater than for the use-all-data method. Confidence in rate estimates is related to tree size, and most clades (70%) analyzed by MRW2018 contained five or fewer species.

## 2. Is the *arithmetic mean* a flawed estimator of central tendency?

I now provide an example of why the comparison advocated by MW2017 and MRW2018 is not valid, by illustrating how this approach (Fig. 1) implies that the arithmetic mean is flawed estimator of the central tendency of a population. The arithmetic mean – arguably the most widely-used statistical estimator in the sciences – is trivial to compute: given a set of numerical observations, we take their sum and divide the result by the number of observations. However, the arithmetic mean is not the only possible (or even best) measure of the central tendency of a particular population from which samples have been drawn. As alternatives, one might use the median, the mode, the geometric mean, or the harmonic mean. Following the statistical test outlined in MW2017 and MRW2018, we can perform a simple simulation exercise to demonstrate that the arithmetic mean is “flawed” and should not be used. The simulation algorithm can be described as follows:

1. We sample a set of N observations from a normal distribution with a specified mean and variance. These N observations represent the complete data and represent all the information we have available for inference about the true population mean.
2. We subsample a set of 4 data points from this larger set of observations, and compute the arithmetic mean of this reduced sample. This approach represents the discard-data method. If 4 observations appear to be insufficient for estimation, again note that 50% of subsampled populations (genera) in the MRW2018 analyses contained 3 or fewer observations. Formally, this approach attempts to estimate *E*(*θ*_*X*_*| X*), where *θ*_*X*_ is the mean for the subsample of these 4 individuals, conditional on the data *X* (4 observations).
3. We then compute the arithmetic mean for the full set of observations. This value is the use-all-data estimate, which is an estimate of *E*(*θ*_*X*_*| X,Y* : *X ∈ Y*). Because the inference model assumes that there is but a single population, the marginal expectation for the set of four individuals sampled above (*X*) is the mean value across the full sample of N individuals (*X,Y* : *X ∈ Y*).
4. We compare point estimates of the discard-data and use-all-data estimates. If they are uncorrelated, we conclude that the arithmetic mean is a flawed estimator and should not be used.

In my simulations, I allowed the true population mean to vary by a factor of five (500%) among replicates, to avoid the criticism that range restriction in the true values accounts for any possible lack of correlation. I performed 13 such paired simulations, in each case drawing the true population mean from a uniform (0.1, 0.5) distribution; *N* = 63 observations for each population were then drawn from a normal distribution with this mean and unit variance. A total of 13 such comparisons were performed, to match the 13 subsamples considered by MRW2018, and I computed the correlation between these 13 paired (discard-data, use-all-data) point estimates.

By seeding the default random number generator in R with a value of 1, we find a striking lack of correlation between arithmetic means computed at these levels (Figure 2). The correlation is a mere *r* = 0.09 (*p* = 0.77). Following MRW2018, we note that: *“These* [numbers] *should be very similar, if not identical. If they are not similar, then we know that at least one of these estimates must be incorrect, even if we do not know which one.”* Some of the numbers in this example are strikingly different: for example, the first estimate in the use-all-data simulation is 0.14, but the corresponding discard-data estimate is 0.35. The greatest disparity overall is the 8th simulated pair, where the use-all-data mean is 0.40, but the discard-data value is *−*0.51. The discard-data estimates thus consistently disagree with the use-all-data values. For this example, the correlation between true values (which are known in this case, as we specified them) and the use-all-data estimates is nonetheless strong (*r* = 0.79, *p* = 0.001), so it is not a lack of information in the data that drives the lack of correlation between use-all-data and discard-data estimates.

**Figure 2.**
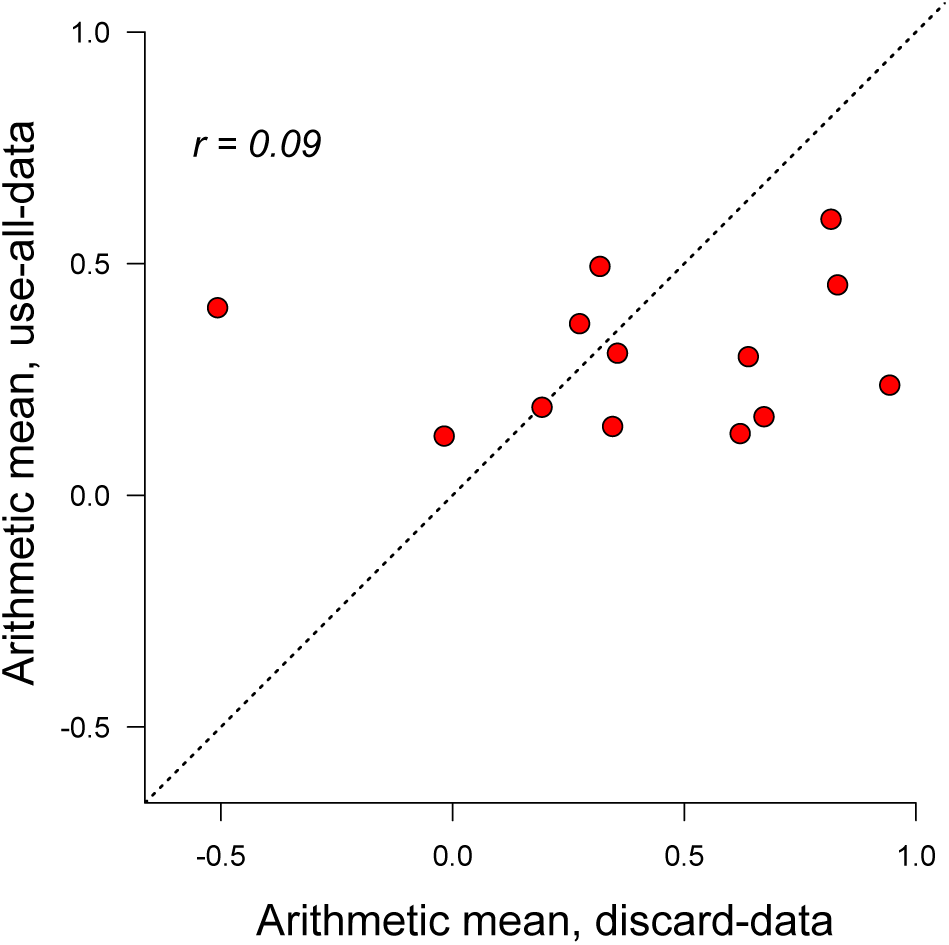
The approach of MRW2018 implies failure of the arithmetic mean as an estimator of central tendency. Arithmetic means were estimated with the discard-data and use-all-data procedure of MRW2018 across a set of 13 populations with different (true) means. Consistent with MRW2018, subgroup size was 4 individuals per sample for the discard-data analysis; a total of 13 populations were chosen for consistency with the MRW2018 tortoise dataset. There is no significant correlation between arithmetic means estimated with the discard-data and use-all-data approach. Under the testing scheme advocated by MRW2018, we would reject the arithmetic mean as a valid inference tool. To replicate this example, see R code available at (src/variance/src/1.arithmetic-mean.R).

We can expand upon these analyses and perform 1000 replicates of the procedure outlined above: in each case, 13 use-all-data estimates are paired with a corresponding discard-data estimate. The correlations and associated p-values are tabulated from each replicate. Surprisingly, 12.3% of such simulations show correlation coefficients that are less than zero (e.g., a negative correlation), and only 19.8% of the correlations are significant (*p <* 0.05). Fully 72.3% show correlations that are less than that observed by MRW2018 for the tortoise dataset (*r* = 0.49). As MRW2018 would conclude: “*Overall, these results show that* [the arithmetic mean] *must be giving estimates that are incorrect in the real world: the two different sets of* [numbers] *are very different, yet both are from* [the arithmetic mean]. *We think that any future studies that use* [the arithmetic mean] *should demonstrate that similar problems do not apply to their analyses.*”

R code to reproduce these results is available at: (src/variance/src/1.arithmetic-mean.R).

## 3. The arithmetic mean is not flawed

The preceding exercise should be recognized for the statistical nonsense that it is. When a test is constructed in this fashion, it is obvious that the variance in the estimators (arithmetic mean) is likely to differ at the two levels of analysis, due to the disparity in sample size alone. For the use-all-data analysis, the arithmetic mean is computed from a sample of 63 observations; the discard-data analysis computes the arithmetic mean from a subsample of just 4 observations. Indeed, we can compute the theoretical variance in the estimators at these two levels of analysis, given these sample sizes. For normal distributions, the variance in the mean of a sample is given by *s*^2^ = *σ* ^2^*/n*, where *σ* ^2^ is the true population variance.

Figure 3 depicts the variance in the arithmetic mean estimator as a function of the sample size. Clearly, the expected variance (e.g., “noise”) associated with estimation is expected to be vastly greater for the small discard-data sample. In this example, the expected variance associated with the arithmetic mean as estimated from the discard-data analysis is 3.02; for the use-all-data analysis, the variance is 0.14. The ratio of variances at these levels of analysis is 21.4. This ratio might seem implausibly large, but as I show below (section 5), the ratio of variances for use-all-data and discard-data analyses in MRW2018 is of a similar or greater magnitude

**Figure 3.**
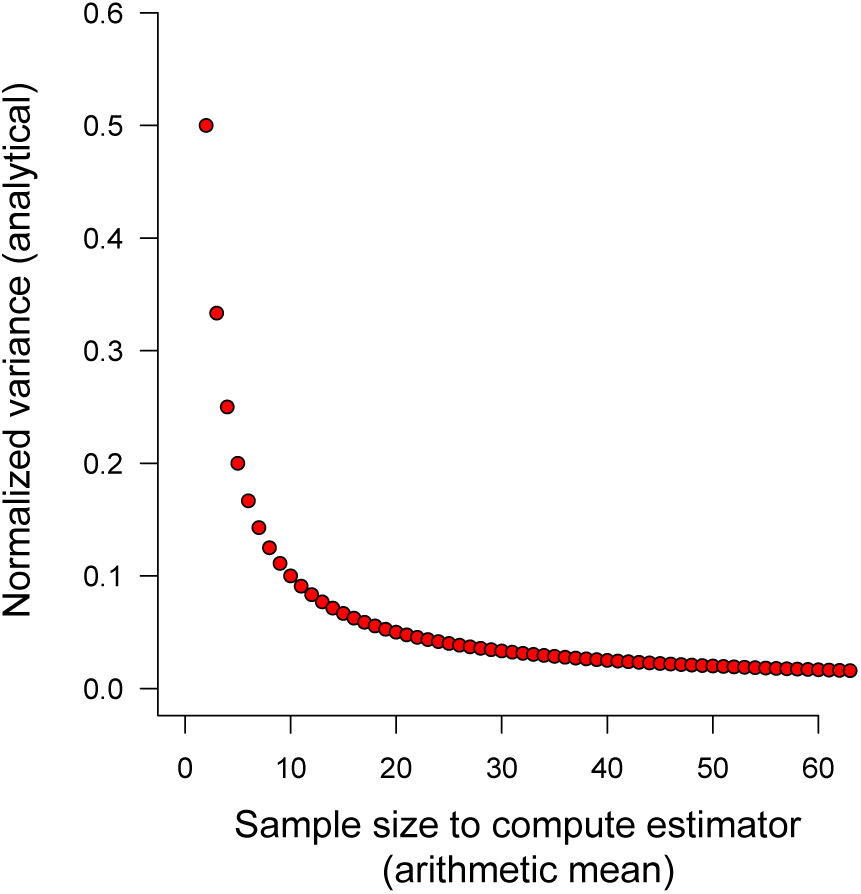
Normalized variance in the arithmetic mean of a sample, as a function of the sample size (number of observations; *N*). Estimation error is inversely related to the sample size and is especially severe for samples with 10 or fewer observations. For these reasons, comparisons between parameter estimates performed for datasets that differ dramatically in the amount of data must be attentive to the effects of sampling variation, particularly if some groups are characterized by few observations. In MRW2018, an estimation procedure was applied to a dataset (tortoises) where 12 of 13 experimental groups contained fewer than 10 observations (species), and where half of all groups contained just 2-3 observations. Y-axis values (normalized variance) are given in units of the true population variance.

In summary, the analyses here provide no information as to whether the arithmetic mean is a flawed measure of central tendency. The lack of correlation illustrated in Fig. 2 is the fully predictable outcome of a comparison between numerical point estimates that ignore sampling variation. R code to reproduce these results is available at (src/variance/src/1.arithmetic-mean.R)

## 4. MRW2018 results are fully explained by sampling variation

The discard-data analyses performed by MRW2018 involve the analysis of very small clades, many of which contain 5 or fewer species. We expect clades with 5 or fewer species to have little biological signal (Rabosky et al 2017), and the resulting posterior distributions of evolutionary rates are expected to have high variance. I perform two exercises below. First, I illustrate - using a phylogeny simulated under known parameters - the consequences of sampling variation for estimates at use-all-data and discard-data levels. Then, using the actual dataset from MRW2018, I perform a power analysis to ask whether the authors would have had power to detect a true correlation given the observed levels of sampling variation in their data.

As a simple illustration of the severity of sampling variation for the inferences in MRW2018, I simulated a phylogeny with 50 taxa under a simple diversification model (speciation rate = 0.1, extinction = 0), using the diversitree package (FitzJohn 2012) and seeding the default random number generator in R with a value of 1. Code to reproduce this example is available at (src/variance/src/2.simple-tree.R).

As in MRW2018, I broke the phylogeny into a set of 12 monophyletic “genera”, each of which contained two or more species. The number of species per genus ranged from 2 to 8, and the median number of tips per genus was 3 (versus 3 in MRW2018). For the use-all-data analysis, I analyzed the complete phylogeny with BAMM, and then estimated the marginal posterior distributions of evolutionary rates for each of the “genera”, or *E*(*θ*_*X*_*| X,Y* : *X ∈Y*). Thus, I estimated a rate distribution for each genus separately, but the rate distribution is conditioned on the complete phylogenetic tree. I then performed discard-data analyses by analyzing each of the 12 genera separately with BAMM, thus discarding all information from the tree more generally and using only the “genus” phylogeny alone (see Fig. 1: discard-data). This latter procedure estimates the quantity *E*(*θ*_*X*_ *|X*).

Figure 4 illustrates the difference between use-all-data and discard-data estimates. The reference phylogeny and four focal genera is shown on the left; corresponding marginal rate distributions from use-all-data and discard-data methods are shown as histograms. For the use-all-data analysis, the BAMM estimated rates (solid blue lines) converge on the true rate with high accuracy. For the discard-data analysis, sampling variation is extreme, to the point that we should ascribe very little confidence to mean rates estimated from small clades. Importantly, the true rate is identical for all clades in Fig. 4, yet the point estimates for each clade under the discard-data approach vary substantially. The tiny clades in Fig. 4 are fully consistent with those analyzed by MRW2018 (e.g., tortoise dataset: median clade size, 3 tips).

**Figure 4.**
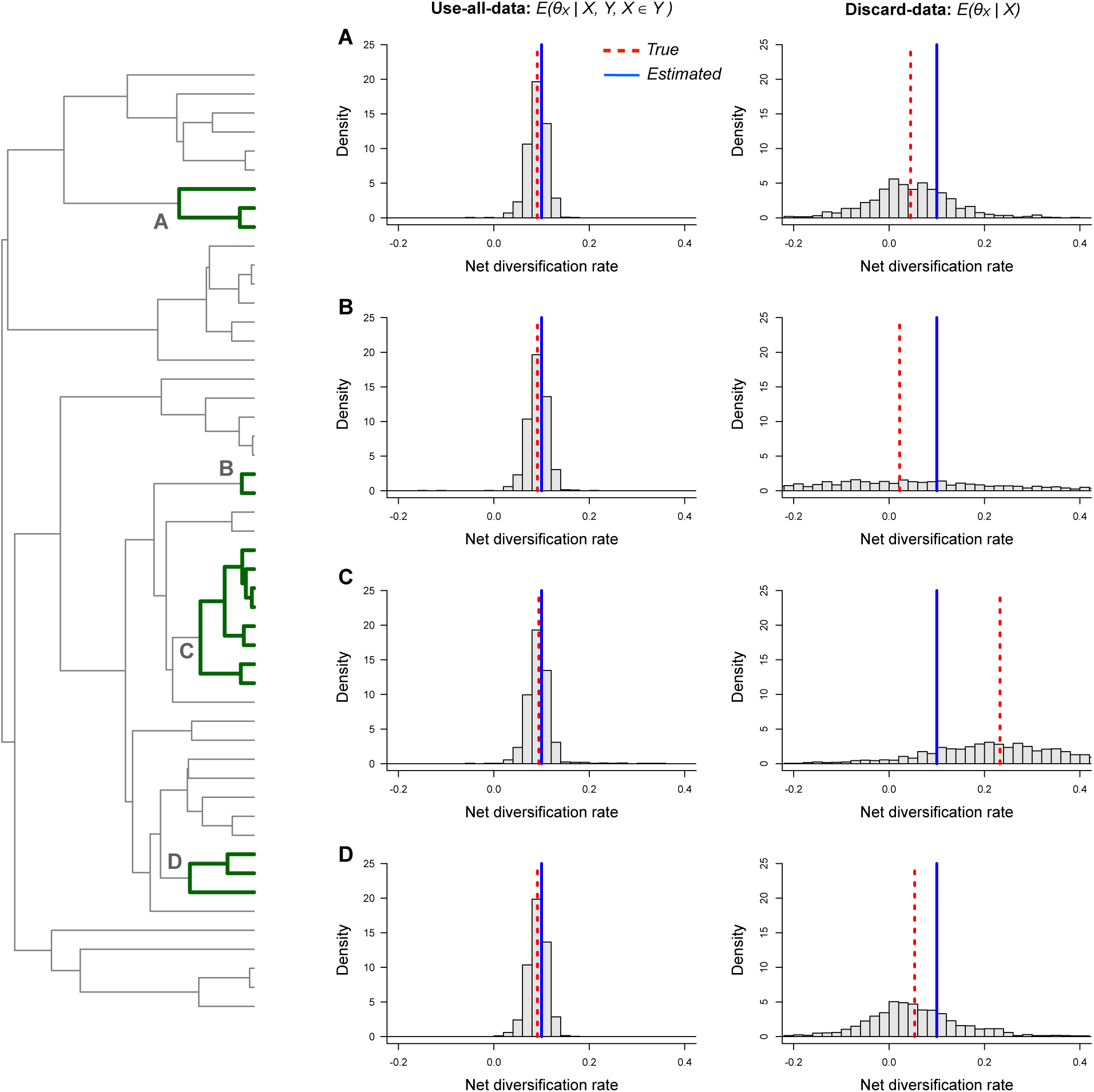
Posterior distributions of evolutionary rates inferred with BAMM for subclades (A, B, C, D) of a phylogenetic tree, where the estimates for each clade are conditioned on the full dataset (use-all-data; left column) and the subclade data alone (discard-data; right column). Each clade thus has two rate estimates: one rate conditioned on all of the data, and another obtained by discarding everything but the focal clade itself. Dashed red line shows true rate for the subclade, and blue line indicates the mean of the marginal posterior rate distribution as inferred with BAMM. The variance of the rate distributions is massively inflated when subclades are analyzed alone, and the corresponding mean rates are much less accurate overall. Subclade sizes illustrated above are commensurate with those analyzed with BAMM by MRW2018. To replicate, see R code available at (src/variance/src/2.simple-tree.R).

For the power analysis, I repeated both use-all-data and discard-data BAMM analyses across the tortoise dataset; these results were not provided by MRW2018, so it was necessary to repeat them. As in Fig. 4, I extracted the marginal evolutionary rates for each genus, first from a full BAMM analysis of the complete tree (use-all-data), and then from a targeted analysis that included only the focal genus (discard-data). Fig. 5 illustrates the marginal posterior distributions of net diversification rates across each of 13 tortoise clades analyzed by MRW2018; the use-all-data and discard-data results are shown in blue and red, respectively. It is immediately clear that the variance in these distributions is much greater in the discard-data analyses, relative to the use-all-data analyses. In Fig. 6, I show the 95% credible intervals on the distribution of rates for each clade under the two analysis procedures. Credible intervals are much larger for the discard-data method (red), relative to the use-all-data method (blue).

**Figure 5.**
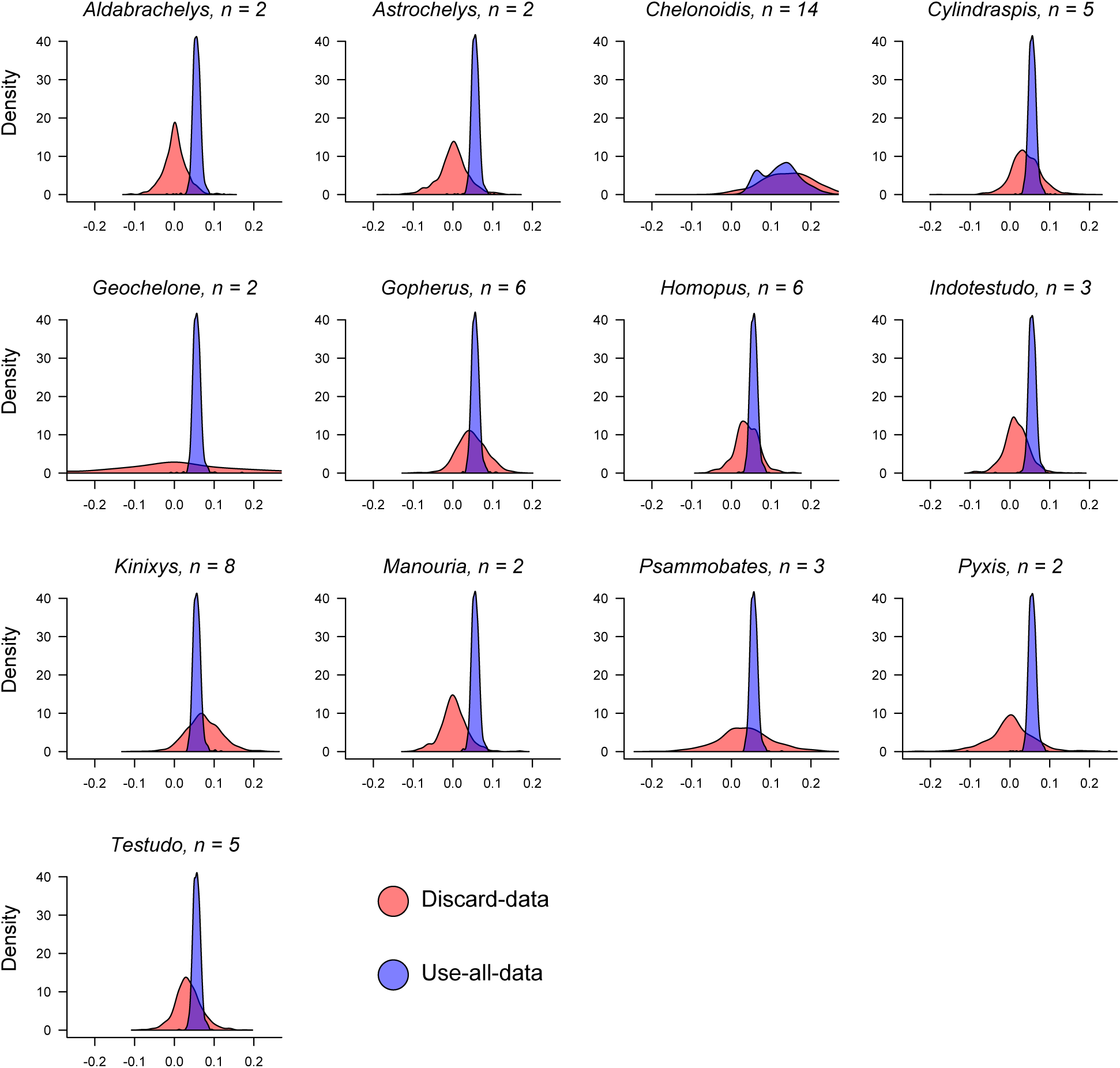
Posterior distributions of evolutionary rates for 13 tortoise subclades, under the discard data and use-all-data approaches. The number of species in the analysis is given next to the genus name, above each density plot. The median number of species in the analyses is 3. The discard-data analyses are associated with much greater sampling variation: the variance of the estimators is, on average, 55 times larger in the discard-data analysis relative to the use-all-data analysis. R code to reproduce this analysis is available at (src/variance/src/3.tortoise-bamm.R).

**Figure 6.**
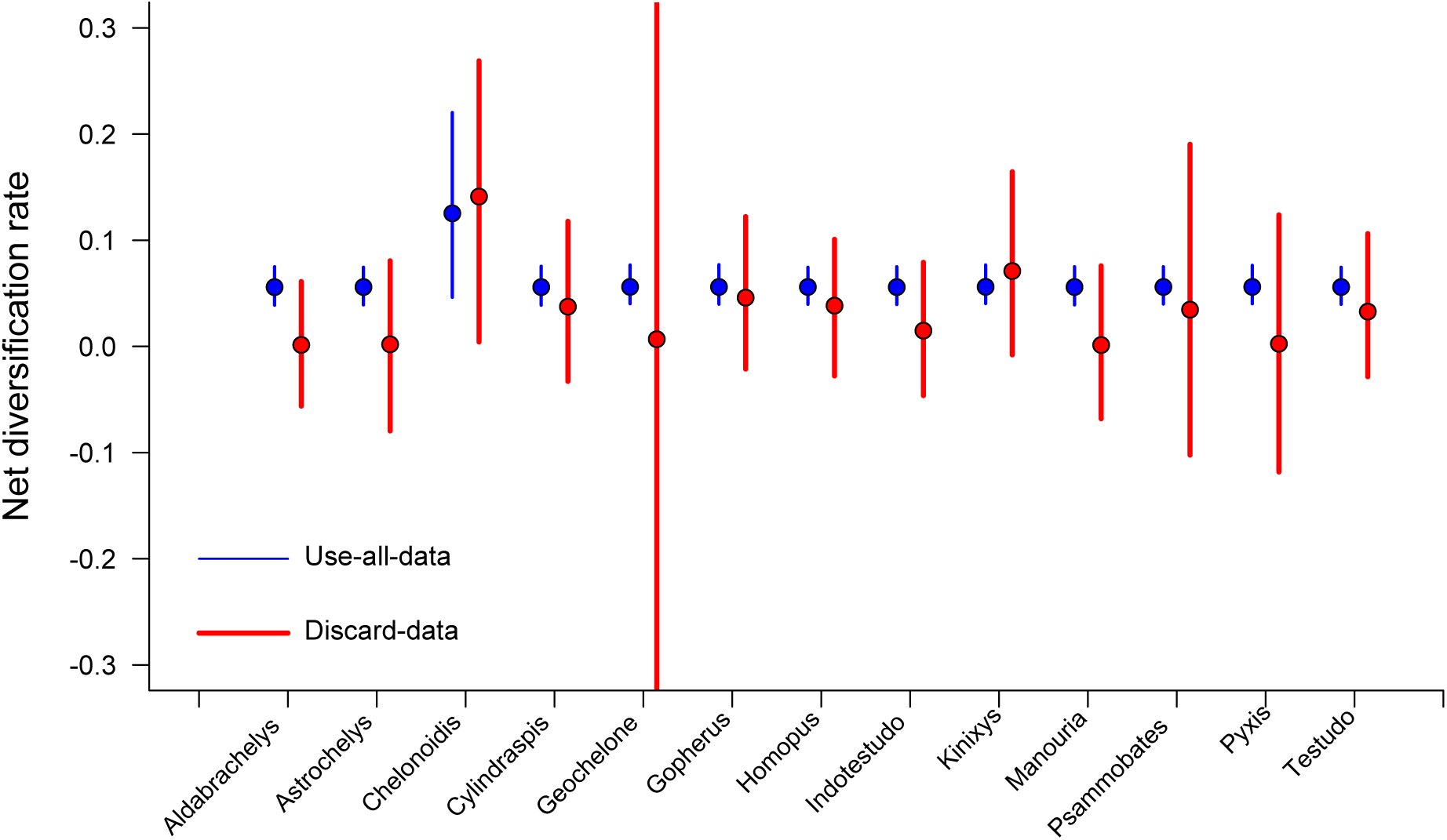
Credible intervals (95%) and medians from the marginal posterior distributions of net diversification rates as estimated for 13 tortoise genera with BAMM. Credible intervals are much larger for analyses conducted with the discard-data scheme, reflecting the small clade sizes analyzed by MRW2018.

I computed the ratio of variances in the inferred posterior distributions for each genus. For example, for the genus Astrochelys (*n* = 2 taxa), the variance in the posterior distribution of rates is 0.0015 from the discard-data method, versus 0.000093 from the use-all-data method; there is thus a 16-fold inflation in variance of rates estimated for this genus with the discard-data approach, relative to the use-all-data approach. Across all 13 clades, the mean variance inflation is 55.2, and the median variance inflation is 15.9. These numbers should not be surprising: because most clades in MRW2018 contain few species, the corresponding estimates at the discard-data level will contain little biological information and the resulting marginal posterior distributions should primarily reflect sampling variation (noise); potential effects of rate priors will also be amplified for very small datasets.

These numbers imply a large impact of sampling variation on the overall results of MRW2018. We now consider precisely how bad the impact could have been, given the overall MRW2018 sampling design. Let us imagine that the use-alldata approach has estimated the true rates for each tortoise subclade with perfect accuracy. Given these rates, we can simulate samples under the discard-data approach, using observed (empirically-estimated) sampling variation. We will assume that the discard-data method is also perfectly accurate and unbiased, but nonetheless contains additional sampling variation. For example, consider the genus Astrochelys: the mean rate from the use-all-data analysis is 0.056, and the discard-data variance is 0.0015. A simulated instance of the discard-data analysis would draw an observation from a normal distribution with these parameters. If the sampling variation is zero, we will observe a perfect correlation between two sets of rates generated in this fashion. The only source of variation between the use-all-data and discard-data estimates is thus the sampling variation, for which the empirically-observed values are used.

Fig. 7 illustrates the distribution of correlation coefficients and p-values that are obtained from the empirical estimates of sampling variation, given a perfect (true) correlation between estimators at the use-all-data and discard-data levels. Across 1000 datasets simulated in this fashion, we find that 74% of simulations show correlations that are weaker than correlation reported by MRW2018 (*r* < 0.49); fully 17.1% of such datasets show *r* < 0 (negative correlations), and only 20.6% of datasets show correlations that are significantly different from zero (*p <* 0.05). Hence, the results from MRW2018 are fully consistent with the hypothesis that the effect they report is fully attributable to neglected sampling variation.

**Figure 7.**
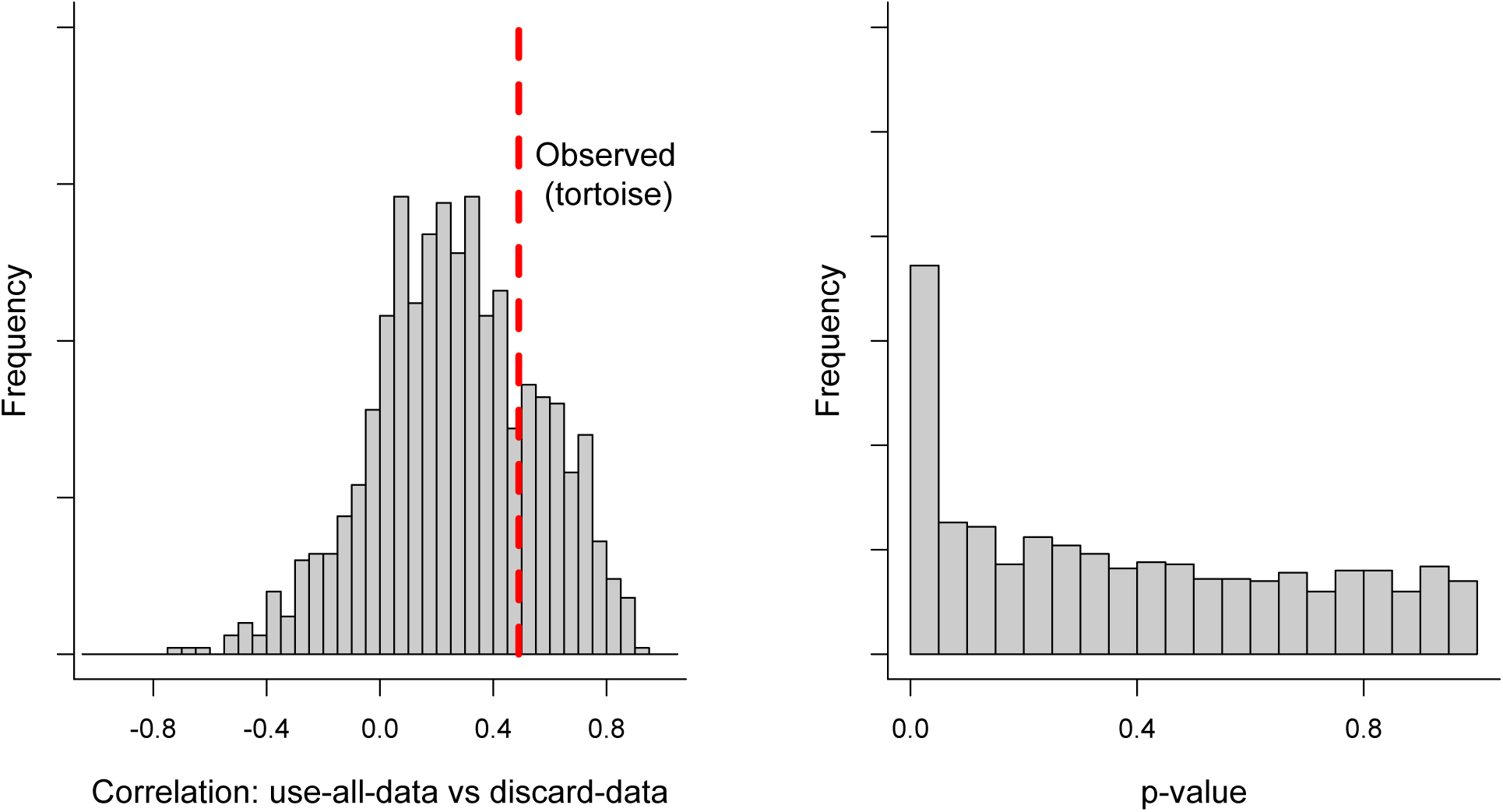
Expected correlation between evolutionary rates estimated at *use-all-data* and *discard-data* levels for the tortoise dataset under the assumption that BAMM rate estimates are perfectly correlated between these levels. In these simulations, all differences in rates between the *use-all-data* and *discard-data* levels comes from sampling variation, which was empirically parameterized following the analysis of MRW2018. This analysis indicates that, even if BAMM has estimated diversification rates with perfect accuracy at one level, the amount of sampling variation is generally sufficient to weaken or eliminate a correlation in rates between the two levels of analysis. To reproduce this analysis, see R code at (src/3.tortoise-bamm.R).

What have we learned from this exercise? Even if BAMM provides unbiased estimates of diversification rates at both levels of analysis, the amount of sampling variation at the discard-data level would be sufficient to erase most of the signal in the data. Given the modest samples of clades (tortoises: 13 clades; tanagers: 30 clades) in the MRW2018 dataset and the high sampling variation in the discard-data estimates, there is no reason to expect significance (*p <* 0.05) of a correlation coefficient computed between rates at use-all-data and discard-data scales.

R code to reproduce the power analysis is available at (src/3.tortoise-bamm.R). All BAMM runs are available in full at (src/variance/data).

## 5. What if there is no variation in diversification rates across the phylogeny?

The approach of MW2017 and MRW2018 will also fail if there is no variation in diversification rates across the focal phylogeny, as pointed out by Rabosky (2018). If the use-alldata and discard-data methods are perfectly unbiased estimators of the true diversification rate, but the diversification rate does not vary, then all variation between the estimators will reflect sampling variation, and the MW2017 and MRW2018 approach will yield an expected correlation of zero. A correlation of zero will be obtained even when the rates are estimated with near-perfect accuracy at all levels, provided that there is some small amount of sampling variation. As Fig. 4 illustrates, evolutionary rates estimated for different genera from a phylogeny that has one true rate can differ substantially due to sampling variation, just as independent samples from any population may differ.

For example, imagine that we have a single phylogeny with 4 clades; the true rates are constant across the phylogeny (e.g., rate = 1). Suppose that the use-all-data approach estimates rates of A = 1.01, B = 0.99, C = 1.01, and D = 0.99; let the corresponding discard-data rates be A = 0.99, B = 1.01, C = 0.99, and D = 1.01. In both cases, the rates have been estimated with extremely high accuracy (mean absolute error = 0.01; 1% of the true value). However, the rates nonetheless show a perfect negative correlation (*r* = *−*1). This is a grave concern in any situation where point estimates are compared across populations with no attention to sampling variation in the point estimates. Because MRW2018 apply their test to empirical trees only, they can offer no assessment of accuracy.

## 6. The misuse of correlation, part I: theoretical insufficiency

The correlation coefficients computed by the authors provide no information about whether BAMM is estimating rates accurately or inaccurately. The correlation coefficient measures the association between two sets of numbers: it is not informative about their absolute accuracy. High correlations between use-all-data and discard-data estimates can be wildly inaccurate, and low correlations are fully consistent with high accuracy (Fig. 8). First, the correlation between two sets of estimates can be zero even if the use-all-data estimates have near-perfect accuracy (Fig. 8, top row); this scenario is fully consistent with the results of MRW2018, given the extreme sampling variation of their discard-data estimators. Second, the correlation between two sets of estimates, and between those estimates and the true values, can be zero even when rates are estimated with high accuracy (Fig. 8, middle). Consider how the correlation coefficient will fare when there is no variation in evolutionary rates across the phylogeny: there is one true rate, but rates for each subclade will be estimated with some amount of error due to sampling variation. The expected correlation in this case is zero, assuming that errors are uncorrelated between use-all-data and discard-data levels. Even if rate variation exists, the extent to which a “significant” correlation can be recovered is a strict function of the variance in the distributions of use-all-data and discard-data estimates. With high sampling variation, even a strong association between rates at these levels will be obscured. Finally, a strong positive correlation between estimates tells us nothing about the overall accuracy of the estimates (Fig. 8, bottom row). This situation may seem unlikely, but can arise in any context where the estimators have correlated errors.

**Figure 8.**
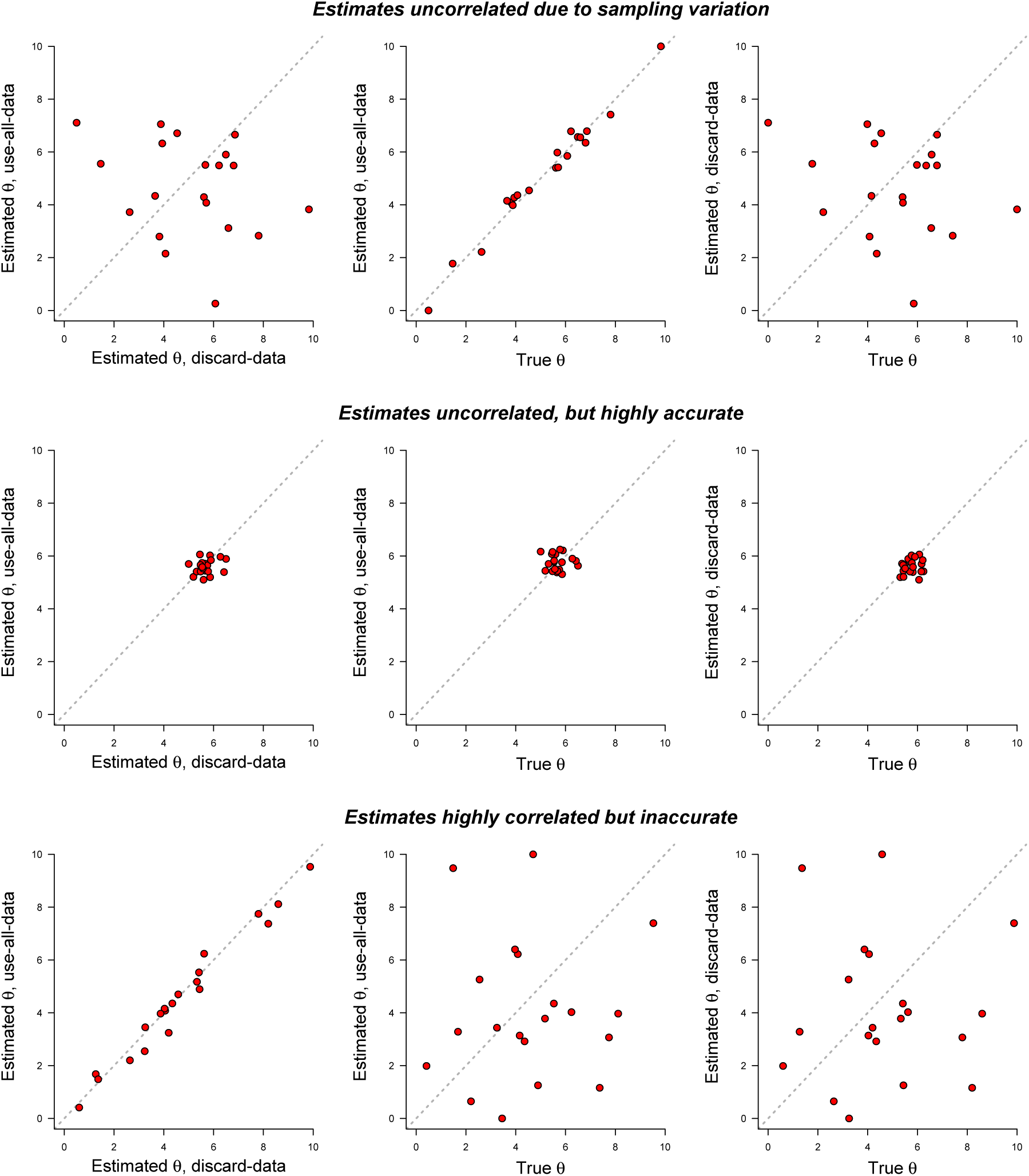
The correlation between two estimates of a parameter *θ* at different hierarchical levels of analysis provides no information about the accuracy of the estimation procedure. Top row: estimated parameters at *use-all-data* and *discard-data* levels are uncorrelated, but the use-all-data estimates are highly accurate. This is the expected outcome when sampling variation confounds estimates at one level of analysis, as in MRW2018. Middle row: No significant correlations are observed in any pairwise comparison, yet both estimates are highly accurate in an absolute sense. Bottom row: estimates from use-all-data and discard-data levels are highly correlated with each other, yet both estimates are uncorrelated with the true rates. This latter situation can arise whenever estimation error is correlated across levels. For example, the implementation of an estimator in a particular software program might be incorrect, leading to parameter estimates that are simultaneously correlated across levels (use-all-data, discard-data) yet uncorrelated with the true parameter values.

## 7. The misuse of correlation, part II: assumptions violated

On top of the issues in Section 6, which invalidate the MW2017 and MRW2018 correlation test in general, the authors violate nearly every statistical assumption associated with this measure of association. I am not arguing here that the authors should have simply used a Spearman correlation (see preceding section), but there are a number of elementary errors in their application of the Pearson correlation. First, the analyses are contaminated by outliers, which invalidate both the computed coefficients and the corresponding p-values. For the tortoise dataset, with just 13 data points, the estimated net diversification rate for a single genus (*Chelonoidis*) is 40 standard deviations outside of the distribution for the remaining 12 observations. For tanagers, the genus *Geospiza* has a rate that is 16.3 standard deviations beyond those of the remaining 29 data points. Second, there is nothing linear about the relationships between the use-all-data and discard-data estimates; such linearity is also assumed by the Pearson correlation. Both of these violations are immediately obvious from visual inspection of Figure 1 from MRW2018.

Third, the Pearson correlation purports to measure the association between independent samples from populations with respective standard deviations *σ*_*X*_ and *σ*_*Y*_. In MRW2018, each clade in the discard-data analysis is drawn from a different error distribution (due to differences in clade size alone), and the empirically-estimated variances from the posterior distributions for individual genera range from 0.00085 to 0.053. This range represents a 62-fold difference in variance across samples that are assumed to have an identical error distribution.

Perhaps the most significant error in the correlation analysis is that the correlation as performed by MW2017 and MRW2018 fails to account for phylogenetic structure in the distribution of diversification rates. The authors assume that the rate estimates for individual genera are independent samples from a particular population. The use-all-data estimates compared by MRW2018 violate this assumption in the extreme. For tortoises, the rate estimates are drawn from a posterior distribution that generally favors just a single rate shift. This one shift, which typically is recovered at the base of the genus *Chelonoidis*, means that BAMM treats the data as though there is an ancestral “slow” rate across most of the tree, which shifted to a “fast” rate along a single branch (leading to *Chelonoidis*). The inference model itself thus imposes strong non-independence on 12 of 13 “observations” for this dataset. In other words, the authors first used a Bayesian model to identify independent groups in their data (BAMM); BAMM then determined that 12 of the genera are not statistically independent, because they share, through their evolutionary common ancestry (c.f. Felsenstein 1985), a common underlying (and inherited) diversification rate. The program then estimates what is effectively a single rate distribution for this set of taxa, and these rates are highly autocorrelated by the inference procedure itself. The rates for 12 taxa thus represent a single data point (sample); they are not independent samples from a single distribution. In the use-all-data analysis, these 12 clades differ in estimated (posterior mean) net diversification rate by a maximum of 0.0026 lineages/my (mean difference: 0.0007) and have an overall coefficient of variation of just 0.04. These numbers are well within the range of variation that could be produced by repeated analyses of the same dataset with BAMM or any other MCMC algorithm. Thus, the authors perform a Pearson correlation with an effective size of just two independent samples.

This latter violation is no different than failing to account for phylogenetic non-independence in any other context. A diversification “rate class”, as estimated by BAMM, is functionally the same as any another discrete character that we might consider. In the case of tortoises, the data suggest that just one character transition occurred across the phylogeny: an ancestral “slow” diversification rate shifted to a “fast” rate in one genus (*Chelonoidis*). There is only a single phylogenetically independent character transition in the dataset, yet MRW2018 treat each genus as a fully independent data point. Twelve of these genera are inferred to share exactly the same character state, by virtue of their shared evolutionary history, and consequently, analyses at the use-all-data scale reveal no variation in rates among most clades (see Fig. 1 from MRW2018).

## 8. Inference models are not equivalent

The preceding analyses demonstrate that the entire effect reported by MRW2018 is consistent with neglected sampling variation; MRW2018 do not control for, report, or discuss any aspects of variation associated with their point estimates. However, it should be noted that there are additional technical issues that lead to non-equivalence of the rate estimates as performed by the authors as they apply the use-all-data and discard-data methods. Critically, the authors use a different mathematical model when estimating diversification rates for the discard-data analysis relative to the use-all-data analysis. Specifically, with the discard-data analysis, the likelihood calculations used by the authors are conditioned on survival of the focal subclade to the present. For the use-all-data analyses, there is no specific conditioning of subclade survival to the present; the survival conditioning is only applied at the base of the full phylogenetic tree. Thus, the basic mathematical model used by these authors at the use-all-data scale is not the same as the model used for the discard-data scale. There is thus a major mathematical difference between the inference models applied at these scales, and the impact of this difference can be very large if extinction is inferred to be high (Nee et al. 1994).

Let *P*_*S,Y*_ denote the probability that the clade representing the full dataset, *Y*, has survived to the present to be observed; *P*_*S,X*_ is the corresponding survival probability for subclade *X*. We can include this additional consideration to formally state the fundamental assumption of MW2017 and MRW2018 as

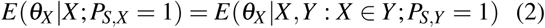

which is clearly invalid in the general sense. The estimators are conditioned on different data, and use a different inference model. This may seem like a trivial difference. It is not. Specifically, the likelihood used by MRW2018 for their discard-data analysis differs from the use-all-data likelihood by the following nonlinear term

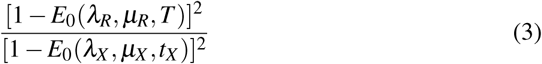

where *E*_0_(*λ,µ, t*) is the probability that a single lineage alive at time *t* with parameters *λ* (speciation) and *µ* (extinction) will go extinct before the present day. Parameters *λ*_*R*_ and *µ*_*R*_ refer to the evolutionary rate parameters at the base (root) of the complete phylogeny, and parameters *λ*_*X*_ and *µ*_*X*_ refer to evolutionary rates at the base of subclade *X*; *t*_*X*_ is the age (from the present) of the basal divergence in subclade *X*, and *T* is the age of the full tree. Thus, the results of the comparisons provided by MRW2018 are uninterpretable, because their assessment is predicated on the assumption that use-alldata and discard-data estimators have the same mathematical expectation. This assumption is not valid: it is unlikely that the set of parameters favored by the model under one conditioning scheme will match the parameter values favored under another conditioning scheme.

This issue is not a “BAMM-specific” problem. Because different inference models were used, we would expect to observe numerical discrepancies between use-all-data and discard-data estimates with any diversification framework, including a simple constant-rate birth-death process. To illustrate, we can show that failure to use the same inference model (due to survival conditioning) appears to explain some of the results from MRW2018. MRW2018 focus in particular on the difference in diversification rate estimates for the genus *Chelonoidis* under the discard-data and use-all-data approaches. The authors report point estimates of rates under the use-all-data and discard-data approaches of 0.125 and 0.058, respectively (again, sampling variation is ignored). However, this magnitude of variation is fully consistent with the difference in inference models used by the authors. To demonstrate this, I used maximum likelihood to fit a constant-rate birthdeath model to the *Chelonoidis* data with and without survival conditioning. With survival conditioning, the estimated rate is 0.04; this accords almost exactly with the discard-data BAMM estimate of 0.06, which performs similar conditioning. Without survival conditioning, the maximum likelihood estimate is 0.18; this increase, relative to the unconditioned analysis, is consistent with the use-all-data BAMM estimate of 0.13, which – like the ML estimate – is not conditioned on survival of this particular genus. Attention to this sort of technical detail, which is of considerable significance in diversification modeling (Nee et al 1994; Etienne et al 2016), is critically important when making sweeping claims about the validity of methods based on numerical comparisons of point estimates. The analyses described above can be repeated using R code found at (src/variance/src/5.nonequivalence-of-models.R).

## 9. Summary and conclusions

MW2017 and MRW2018 conduct analyses of empirical datasets at two hierarchical levels that use different amounts of data and with different variance structures, which I have referred to as the *use-all-data* and *discard-data* approaches. The authors find that point estimates of evolutionary rate parameters at these two levels of analysis are numerically different, and conclude that the method used for estimation is therefore flawed. These conclusions are profoundly misleading, for multiple reasons (Table 1).

Critically, there is no mathematical expectation that estimates at *use-all-data* and *discard-data* levels should be the same. The estimators they compare are theoretically expected to give different results, on account of both (1) differences in the datasets that underlie their estimation procedure, and (2) differences in the actual inference models themselves. That these estimators are not the same is a trivial mathematical observation that invalidates their empirical analyses on first principles. There is no theoretical reason that estimates at their two levels of analysis are guaranteed to be identical or even similar, even if rates at both levels are estimated with perfect accuracy and estimation error is zero (equation 1; equation 2).

Even more importantly, MW2017 and MRW2018 ignore sampling variation (variance) in their analyses. Because they estimated population parameters from small samples (median: 3 observations per experimental group), the sampling variation associated with their estimators is enormous. MRW2018 stands as perhaps the only study in the literature to apply the method under consideration to experimental groups (phylogenies) with fewer than 20 observations (tips). There is nothing inherently wrong with applying inference methods to extremely small populations, provided that sampling variation and impact of priors (for Bayesian inference) are characterized. However, these factors are ignored entirely by the articles in question. The 55-fold difference in variance between estimators at the use-all-data and discard-data levels of inference is sufficient to erase any meaningful conclusions that can be drawn from the comparisons in these articles (Section 4; Fig. 7).

Variation is perhaps the most general feature of the world around us; it pervades everything and anything that we might wish to measure in nature and forms the basis for evolutionary change itself. Felsenstein (1985) famously said that there is no doing comparative biology without taking phylogenies into account. We might say the same for the role of variance in science more generally.

